# Zebrafish xenografts to isolate unique human breast cancer metastatic cell populations

**DOI:** 10.1101/2021.12.07.471608

**Authors:** Jerry Xiao, Joseph R. McGill, Apsra Nasir, Alexander Lekan, Bailey Johnson, Devan J. Wilkins, Gray W. Pearson, Kandice Tanner, Hani Goodarzi, Eric Glasgow, Richard Schlegel, Seema Agarwal

## Abstract

Cancer metastasis is a critical culprit frequently blamed for treatment failure, drug resistance, poor prognosis, and high mortality rate among all human cancers. Laboratory efforts to isolate metastatic cell populations have typically been confined to mouse models, which are time-consuming and expensive. Here, we present a model system based on xenografting zebrafish embryos to select for cells that are predisposed to progress through the early stages of metastasis. This model requires only 3-5 days to achieve distinct intravasation to the zebrafish circulatory system. The metastatic cells are easily tracked in real-time as they migrate, as well as isolated and propagated *in vitro*. Once expanded, molecular characterization of the serially derived invasive cell populations from the tails of the zebrafish accurately predicts genes, signaling pathways, protein-protein interactions, and differential splicing products that are important for an invasive phenotype. This zebrafish model therefore offers a high-throughput and robust method for identifying gene targets critical for cancer metastasis.

## Introduction

Up to 90% of cancer-related deaths have been attributed to metastasis^1^. Cancer metastasis is frequently associated with treatment failure due to drug resistance, poorer prognosis, and high mortality in all cancers^2^. In breast cancers, the presence of metastasis reduces 5-year expected survival rates to a dismal 28% compared to 99% survival in localized breast cancer^2^. The process by which cancer cells metastasize from a primary node is typically spoken of through the completion of a series of stages: (1) intravasation into the circulatory system; (2) travel through the circulatory system; (3) extravasation through the blood vessel endothelia; and finally (4) dormancy and/or colonization at the metastatic site^1^. Unfortunately, the study of cancer metastasis and identification of therapeutics specifically targeted towards preventing metastasis has been limited by a lack of available models for their study^3, 4^.

Presently, few model systems allow for the easy isolation and study of a metastatic cell population from clinically non-metastatic patients. The first *in vivo* model for metastatic disease was developed in the 1970s by Fidler and Kripke, who injected cultured B16 melanoma cells into mice^5^. Development of *in vivo* metastasis models has since focused on the use of nude, athymic mice, or severe combined immunodeficient (SCID) mice, which lack functional B and T cells ^6^. However, the rodent models suffer from disadvantages that limit their application to understanding the metastatic process or developing drugs that can target the metastatic process. For example, the long latency period for the rodent models (typically 4-6 months) prevents the creation of large-scale, high-throughput drug screens^6^. Additionally, characterization of cells that undergo metastasis is only feasible after euthanasia, providing limited resolution to cells undergoing active metastasis using rodent models^6^. Because of these limitations, rodent models for cancer are typically considered better suited for the study of direct antitumor effects on primary tumors rather than on metastasis^4^. Alternatively, *in vitro* models, such as the transwell invasion assays using either matrix or endothelial cells or various microvessel platforms cannot replicate the crucial multistep process required for the metastatic process^7–9^. Metastasis is an increasingly complex production supported through the inclusion of multiple actors such as tumor-associated macrophages and neutrophils^10–12^, stromal components such as cancer-associated fibroblasts^13, 14^, and even unexpected contributors such as platelets^15^. Furthermore, *in vitro* models also lack the tumor microenvironment, critical for tumor progression that is present in *in vivo* models^8, 16, 17^. As researchers are enlightened to the role that each of these individual players plays in metastasis, *in vitro* models become increasingly obsolete for the accurate evaluation of cancer metastasis^4^. As an alternative, some studies have suggested profiling circulating tumor cells (CTCs). However, methods for evaluating CTCs are exceedingly difficult due to their rare numbers within the circulation^18^.

In recent years, an animal model using the zebrafish (*Danio Rerio)* has emerged as a viable alternative^19^. Zebrafish models of melanoma, prostate, salivary, and breast cancers, show histopathological similarities to their human counterparts when engrafted^20–23^. Furthermore, many of the molecular components involved in metastasis are highly conserved across humans and zebrafish^24–26^. We and others have also shown that cancers modeled in zebrafish display similar chemotherapy sensitivity and therapeutic responses when compared to current rodent-based models^22, 26–28^. In addition to fulfilling critical criteria as an *in vivo* model, zebrafish offer several advantages over other models. For the first two-three weeks, zebrafish have a robust innate immune system but have not yet developed a mature adaptive immune system, therefore requiring minimal genetic manipulation of host animals for successful engraftment of human cancer cells^29^. Furthermore, cells engrafted in 2dpf zebrafish can survive for up to 10 days, and only require the transplantation of a few hundred cells as opposed to millions of cells that need to be transplanted into a single mouse^19, 30^. Furthermore, transgenic zebrafish lines that have been engineered to express fluorescent vasculature can leverage the naturally transparent nature of early zebrafish to allow for non-invasive, real-time imaging of cancer cell development and migration^31, 32^. Additionally, the cost of zebrafish husbandry in a vivarium is significantly less than those required for mice^33^. Finally, each zebrafish breeding pair can yield as many as 300 embryos allowing for rapid and robust scalability of experiments^25, 28^.

Here, we report a model utilizing 2dpf zebrafish embryos that leverage the benefits of zebrafish for the selection of a cancer cell population analogous to human CTCs. Specifically, upon engraftment within the yolk sac of 2dpf zebrafish, a small subpopulation of cells will invade through the endothelium into the zebrafish circulatory system, arresting within the caudal plexus of the zebrafish within 5 days. These cells can subsequently be physically separated from the initial engraftment and propagated *in vitro*. Serial injection of the putative invasive subpopulation through this zebrafish intravasation model subsequently selects for a cell population with distinct phenotypic and transcriptomic differences from its parental population. Notably, the resulting cellular population accurately identifies enrichment of genes and pathways involved in establishing an invasive phenotype. Taken altogether, this model enables the rapid and robust isolation of a population of cancer cells predisposed to intravasation within an *in vivo* context, providing researchers with a powerful tool for the study of the early stages of cancer metastasis.

## Results

### Zebrafish xenografts can be used to differentiate functionally invasive cell populations

We first sought to establish that cell transplantation into 2dpf zebrafish could differentiate between cells that are expected to intravasate and those that are not within a reasonable time frame. To begin, we injected cells from two human epithelial breast cancer cell lines, MCF7 and MDA-MB-231, into the yolk sac of 2dpf zebrafish embryos. In mouse models, the breast adenocarcinoma MCF7 cell line has consistently been shown to be poorly metastatic *in vivo*^34^. On the other hand, MDA-MB-231 cells, which were derived from the pleural effusion of a patient with invasive triple-negative ductal carcinoma, are routinely used as a model for aggressive late-stage cancer^35, 36^. Based on our prior studies, each zebrafish yolk sac could be injected with up to 200 cells to allow for both high-throughput screenings while also maintaining zebrafish viability^22^. Consistent with their behavior in mice, transiently labeled MCF7 cells exhibited minimal intravasation and arrest within the tail of the zebrafish compared to MDA-MB-231 cells after 5 days (**Fig. 1a, b**).

**Figure 1:**
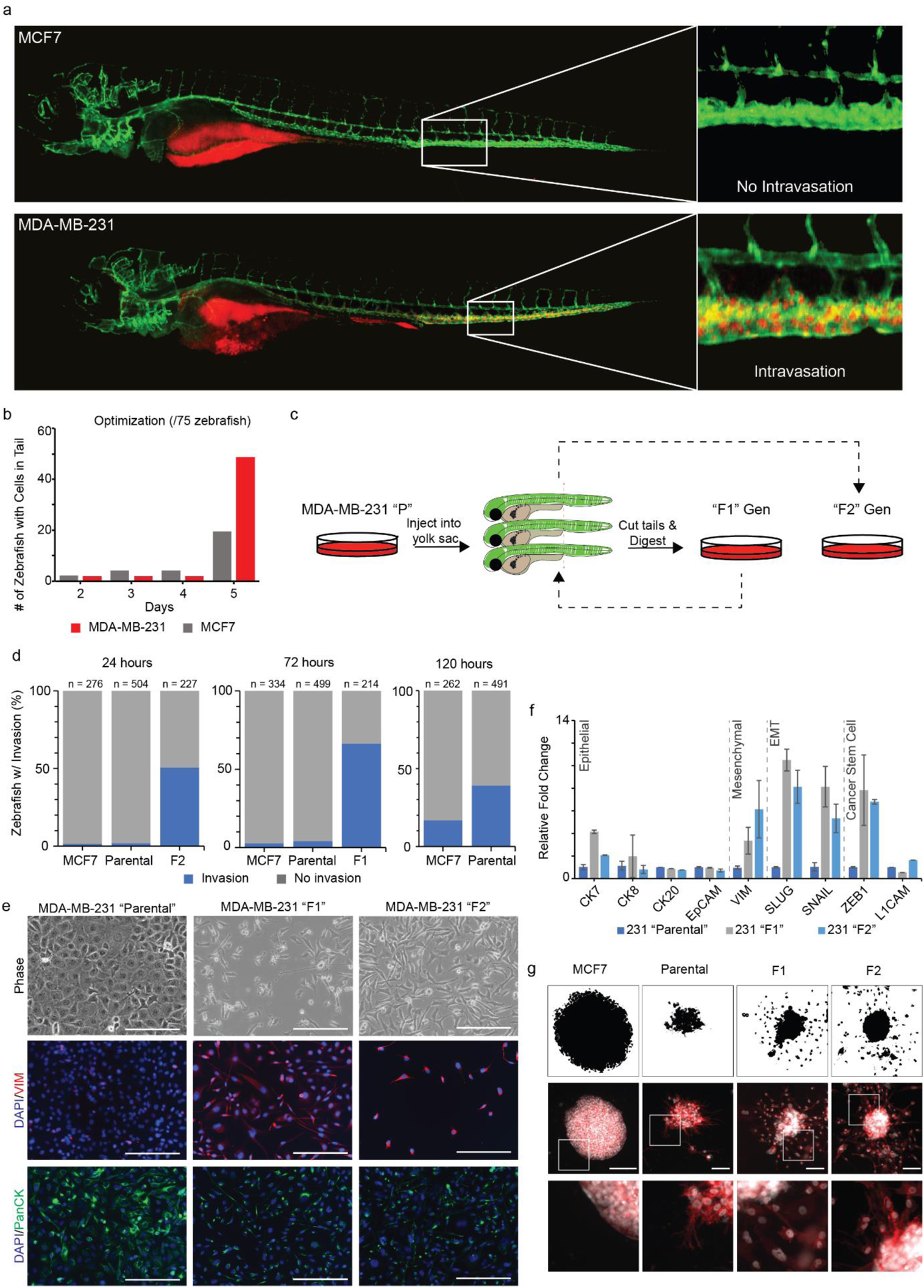
2 days post-fertilized zebrafish embyros can be used to identify differentially invasive cellular populations. (a) Injection into the yolk sac of MCF7 and MDA-MB-231 cells results in cell arrest within the caudal plexus in only MDA-MB-231 injected zebrafish. (b) Cells appear within the tail of the zebrafish within 5 days of injection. (c) Visual workflow of proposed serial transplantation of the MDA-MB-231 heterogeneous parental population, which yields the F1 and F2 subpopulations. (d) 231 parental, F1, and F2 cells have progressively faster time-to-arrest within the zebrafish tail. ≥ 40% of injected zebrafish have cells visibly arrested within the tail in 120, 72, and 24 hours, respectively. (e) Phase and immunofluorescence images of resulting *in vitro* cultured 231 parental, F1, and F2 cells after nearly a year in culture. (f) qRT-PCR amplification of epithelial (CK7, CK8, CK20, EpCAM), mesenchymal (VIM), EMT TFs (SLUG, SNAIL), and cancer stem cell markers (ZEB1, L1CAM) of the three subpopulations. (g) Clusters of MCF7, MDA-MB-231 parental, F1, and F2 cells were embedded within a 3-dimensional Matrigel-based extracellular matrix and allowed to invade over 24 hours. Here, Hoechst stain for cell nuclei is shown in white and Phalloidin is depicted in red. Top = ImageJ mask of cell clusters used for analysis; Middle = phase microscopy, scale bars = 100 µm; Bottom = A 4x digitally generated inset highlighting edges of clusters.

Based on these initial results, we designed a study involving the serial injection of MDA-MB-231 cells into 100 transgenic *flk:gfp* labeled zebrafish (**Fig. 1c**). An initial “parental” population of MDA-MB-231 cells was injected into the yolk sac of zebrafish and monitored for arrest within the tail of zebrafish. When >50% of zebrafish exhibited cells within the caudal plexus, the tails were cut, and any labeled cells were isolated and grown *in vitro* to form the “F1” generation. Once grown, the F1 population was similarly engrafted into the yolk sac of 2dpf zebrafish embryos, subsequently yielding the “F2” population. Final engraftment of the F2 population was then performed to evaluate their invasive phenotype. Upon injection, we observed that the parental, F1, and F2 populations took increasingly shorter amounts of time to achieve >50% intravasation (**Fig. 1d**). While zebrafish injected with parental cells only achieved ~40% intravasation after 120 hours, F1- and F2-injected zebrafish achieved >50% intravasation within 72 and 24 hours, respectively. Overall, both rounds of serial injection to generate the F1 and F2 populations as well as culture growth required ~30 days.

Upon *in vitro* culture, the parental, F1, and F2 populations also exhibited dramatically different morphological phenotypes (**Fig. 1e**). While the parental population exhibited a cobblestone appearance commonly associated with epithelial cells, the F1 and F2 populations exhibited a more protruded, spindle-like morphology associated with invasive, mesenchymal cells ^37^. This epithelial-to-mesenchymal (EMT) phenotype change was also confirmed by increasing vimentin, a mesenchymal marker, and decreasing cytokeratin, an epithelial marker, expression via immunofluorescence (**Fig. 1e**). These trends were similarly confirmed in quantitative reverse-transcriptase PCR (**Fig. 1f**). qRT-PCR of epithelial (CK7, CK8, CK20, EpCAM), mesenchymal (VIM), EMT TFs (SLUG, SNAIL), and cancer stem cell markers (ZEB1, L1CAM) all exhibited trends consistent with an invasive cellular population in the F1 and F2 populations. Notably, the F1 and F2 populations have retained these differences through nearly 2 years of *in vitro* expansion.

Next, we sought to evaluate whether the distinct invasive behaviors of the parental, F1, and F2 generations would be retained through an *in vitro* invasion assay. After one year of culture, MCF7, parental, F1, and F2 were clustered and embedded within a 3-dimensional invasion assay and evaluated for invasive behavior for 24 hours. As expected, MCF7 clusters did not exhibit any invasion in this extracellular matrix. Parental, F1, and F2 clusters exhibited invasion with increasing intensity, consistent with their behavior *in vivo* (**Fig. 1e**). Importantly, while parental clusters did exhibit protrusions after 24 hours, there was minimal cell body migration. By comparison, both the F1 and F2 clusters exhibited significant migration of cell bodies, suggesting the parental population had not fully invaded compared to the F1 and F2 populations.

*Selected F1 and F2 populations accurately enrich for metastasis-associated genes and pathways* In cancer, only a small subset of cells completes the metastatic cascade and result in the formation of a metastatic nodule^5, 38^. Accurate prediction of the particular subpopulation of cells from an initial pool of heterogeneous cancer cells would be a critical tool necessary to support the discovery of metastasis-targeting therapeutics^4^. Therefore, we next sought to determine whether serial transplantation through the zebrafish model could accurately select for a subpopulation of cells that are enriched for metastasis-associated drivers.

To address whether the parental, F1, and F2 populations were transcriptionally distinct, we performed next-generation bulk RNA-sequencing. Principal component analysis of sequenced RNA libraries from the three respective populations revealed significant changes between all populations (**Fig. 2a**). Consistent with our depicted phenotypes, both the F1 and F2 subpopulations expressed lower epithelial (KRT8, KRT18, KRT19, CLDN3, CLDN4, EGFR, and DSP) and higher mesenchymal (VIM, CD44, SNAI2, COL6A3, ITGA5, IL6) genes compared to their parental source (**Fig. 2b**). Overall, using their parental source population as a common control, an absolute log 2-fold change cutoff of at least 1.5, and an adjusted p-value cutoff of 0.05, the F1 and F2 populations were upregulated in 842 genes and downregulated in 875 genes (**Fig. 2c**). Among those genes that were most significantly upregulated were CSF3, G0S2, COL7A1, IL16, SAA2, C15orf48, and DNER.

**Figure 2:**
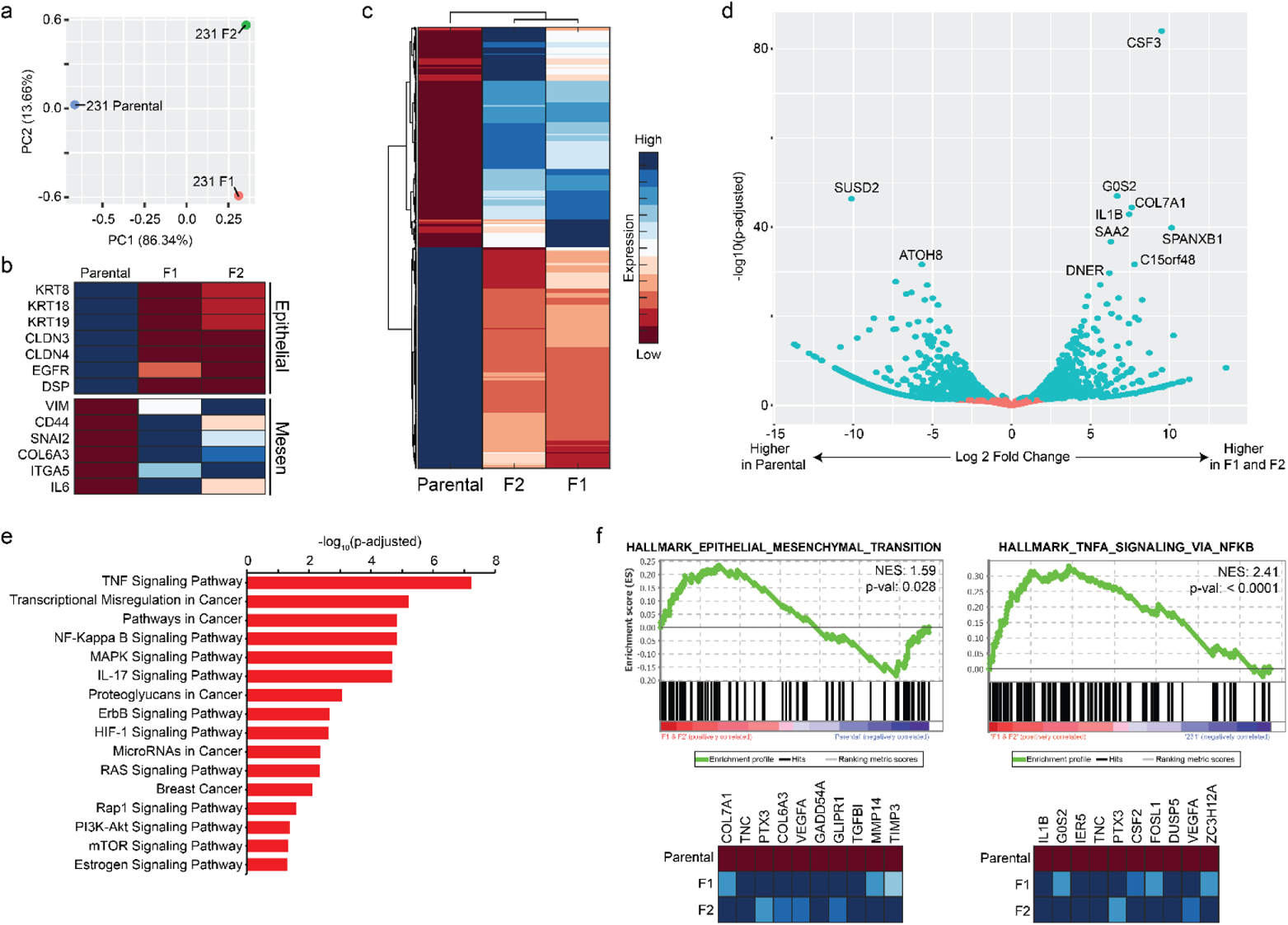
RNA-sequencing of MDA-MB-231 derived parental, F1, and F2 populations was performed. (a) Principal-component analysis reveals transcriptomic distinction between all three subpopulations. (b) A heatmap depicting RNA-sequencing expression of a select panel of epithelial and mesenchymal markers is shown. (c) A clustergram of the overall transcriptomic landscape of the parental, F1, and F2 populations. (d) A volcano plot depicting the most significantly upregulated and downregulated genes when comparing the F1 and F2 populations relative to the common parental control. (e) A KEGG analysis was performed, revealing enrichment of several important cancer-associated pathways in the F1 and F2 populations. (f) GSEA analysis identified enrichment of the hallmarks gene set Epithelial-Mesenchymal Transition and TNFα Signaling via NF-kβ, among other gene sets.

### F1 and F2 populations are enriched for breast cancer-associated pathways and protein interactions

While differential gene analysis provides researchers with an ability to examine trends in individual genes, pathway enrichment analysis provides mechanistic insight based on broader gene sets, leveraging the promises of next-generation sequencing^39^. The Kyoto encyclopedia of genes and genomes (KEGG) is a publicly available database of curated gene sets containing various cancer-associated gene sets as well as common signaling pathways^40^. Using the parental population as a common control, KEGG analysis identified enrichment of several cancer-associated gene sets in the F1 and F2 populations relative to parental such as transcriptional misregulation in cancer (hsa05202), pathways in cancer (hsa05200), and breast cancer (hsa05224) (**Fig. 3e**). In addition to these cancer-specific gene sets, the F1 and F2 populations were also significantly enriched for TNFα, NF-Kβ, MAPK, IL-17, ErbB, HIF-1, RAS, RAP1, PI3K-AKT, and mTOR signaling pathways. Gene Set Enrichment Analysis (GSEA) is another computational method commonly used to determine whether there is a significant difference in pathway expression between biological states^41, 42^. Once again using the parental population as a common control, GSEA analysis identified enrichment of several gene sets in the F1 and F2 populations relative to their parental source, including the hallmark gene set EMT (NES 1.59, p-value = 0.028, **Fig. 2f** and **Table S2**) as well as TNFα signaling, in harmony with KEGG analysis.

**Figure 3:**
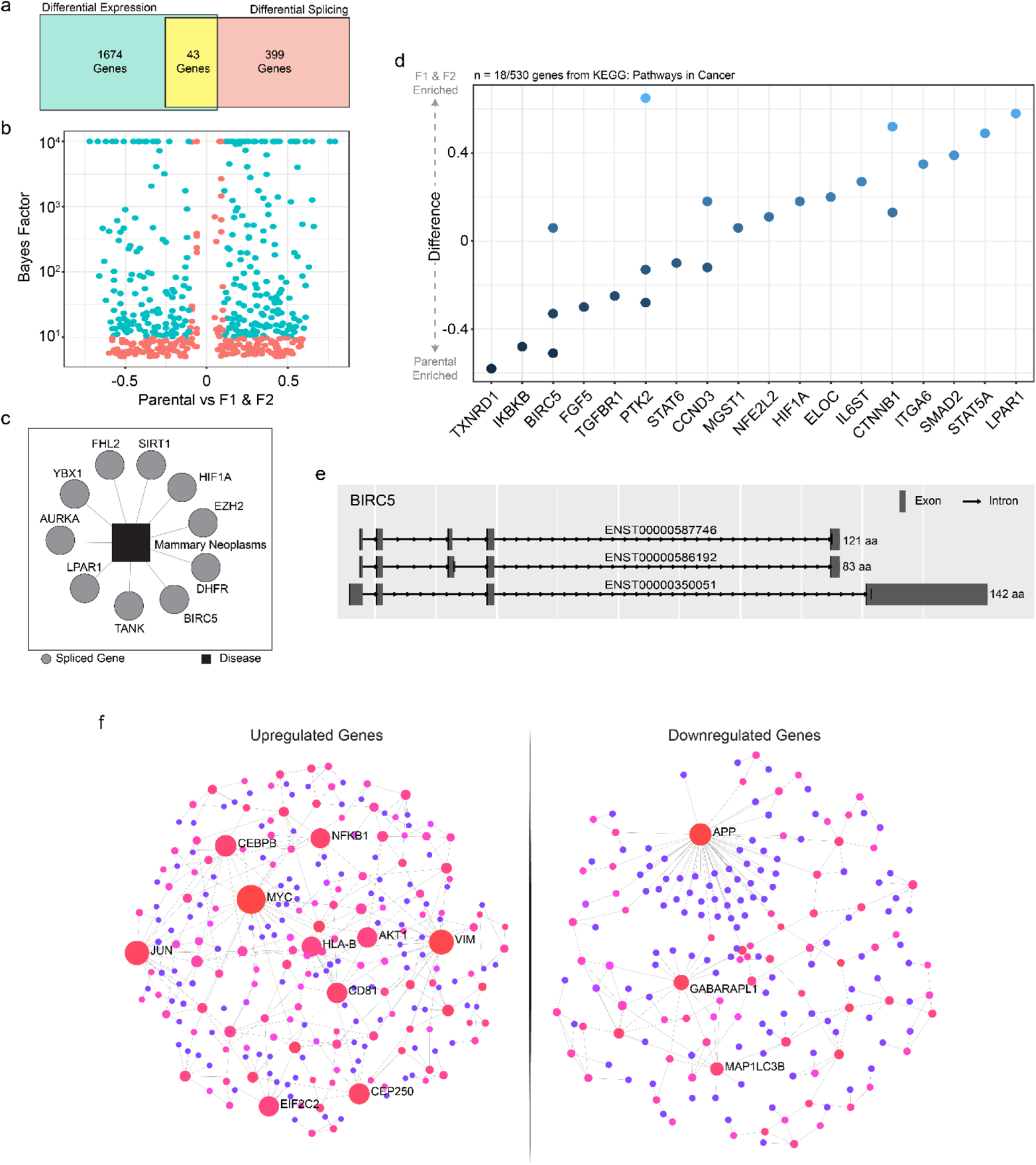
Differential RNA-splicing and protein-protein interactions associated with cancer and metastasis were enriched in the F1 and F2 populations compared to parental population. (a) Overall, 107 genes were both significantly differentially expressed and differentially spliced. (b) A volcano plot of the 526 differentially spliced transcripts. (c) Gene-disease association analysis using DisGeNET revealed differentially spliced genes associated with mammary neoplasms. (d) KEGG pathway analysis identified differentially spliced genes included within the Pathways in Cancer geneset. (e) BIRC5 was one of the few genes that was both differentially expressed and differentially spliced. The three spliced variants of BIRC5 differentially spliced in the F1 and F2 populations are shown mapped. (f) Mammary tissue-specific protein-protein interactions identified the most interconnected proteins involved in the upregulated (left) and downregulated (right) genes.

### The F1 and F2 populations are enriched for RNA splicing events and protein-protein interactions associated with cancer metastasis

While the existence of differential transcriptomic variations linked to metastasis in breast cancer is well studied, the presence of differential RNA splicing events is less explored. Given the limited absolute number of genes encoded by the human genome, it is safe to assume that differential RNA-splicing, which can result in variations of proteins given a similar genomic template, might play a critical role in breast cancer metastasis^43^. Using a computational framework, we identified a total of 526 differentially spliced products spanning 442 unique genes when evaluating the F1 and F2 subpopulations relative to their parental counterparts (**Fig. 3a & 3b**). Overall, 43 genes displayed both differential expression as well as differential splicing. DisGeNET is an integrated platform for evaluating gene-disease associations^44^. When evaluating known gene-disease associations among the 443 unique genes differentially spliced, one of the top associations identified was mammary neoplasms, which were associated with differentially spliced genes such as HIF1A, BIRC5, LPAR1, SiRT1, and others (**Fig. 3c**). Next, KEGG analysis was performed, identifying enrichment of genes belonging to the KEGG gene set Pathways in Cancer (*pval* = 0.0347, **Fig. 3d**). Notably, of the genes identified, BIRC5 aka Survivin is associated with both mammary neoplasms and pathways in cancer. BIRC5 and its splice variants have previously been shown to associate with a poorer overall prognosis in metastatic breast cancer^45, 46^. Clinical studies targeting survivin and its splice variants with surviving antagonists have been proposed with promising results^46^. In this study, we identified differential splicing of three separate variants of BIRC5, each of varying translated protein length (**Fig. 3e**).

Next, we sought to evaluate whether the F1 and F2 populations were enriched for metastasis-associated protein-protein interactions (PPIs)^47^. Recent studies investigating PPIs have given life to the possibility of targeting important interactions to inhibit extracellular signals in cancer^48^. We conducted a network analysis of mammary tissue-specific PPIs within either the up- or down-regulated genes of the F1 and F2 populations (**Fig. 3f**). Among genes that were significantly upregulated in the F1 and F2 populations, MYC, VIM, JUN, CD81, NFKB1, CEBPB, HLA-B, and CEP250 all exhibited high interconnectedness (degree ≥ 10) with other upregulated proteins (**Fig. 3f** and **Table S3**). On the other hand, downregulated proteins APP, GABARAPL1, and MAP1LC3B exhibited high interconnectedness (degree ≥ 10) among the downregulated proteins (**Fig. 3f** and **Table S4**). All-in-all, both differential splicing and PPI network analysis revealed that the F1 and F2 populations were enriched for phenotypes consistent with a metastatic signature.

### The zebrafish model identifies functionally important drivers of cellular invasion

Importantly, one of the key criteria for the selection of the F1 and F2 subpopulations was their ability to successfully invade through the yolk sac and intravasate through the endothelium of the zebrafish circulatory system. We therefore sought to determine whether the increased invasiveness of the F1 and F2 populations could be inhibited through an *in vitro* functional study. We began by identifying seven target genes that spanned a variety of cellular functions and were all upregulated within the F1 and F2 populations. The seven genes identified were SNRPA1, MT1X, SRGN, CTSD, S100A11, SERPINE1, and DDIT4 (**Fig. 4a**). Based on their expression in RNA-seq data, we determined to investigate whether knockdown of these genes within the F2 subpopulation by short hairpin RNA (shRNA) could reduce cellular invasion to levels consistent with their parental source. To verify successful knockdown, we utilized qRT-PCR amplification of these targets in the parental, F1, F2, and F2 infected with a scramble shRNA cell lines. Based on qRT-PCR, expression of the seven target genes was successfully reduced to levels like their parental counterpart in all genes (**Fig. 4b**).

**Figure 4:**
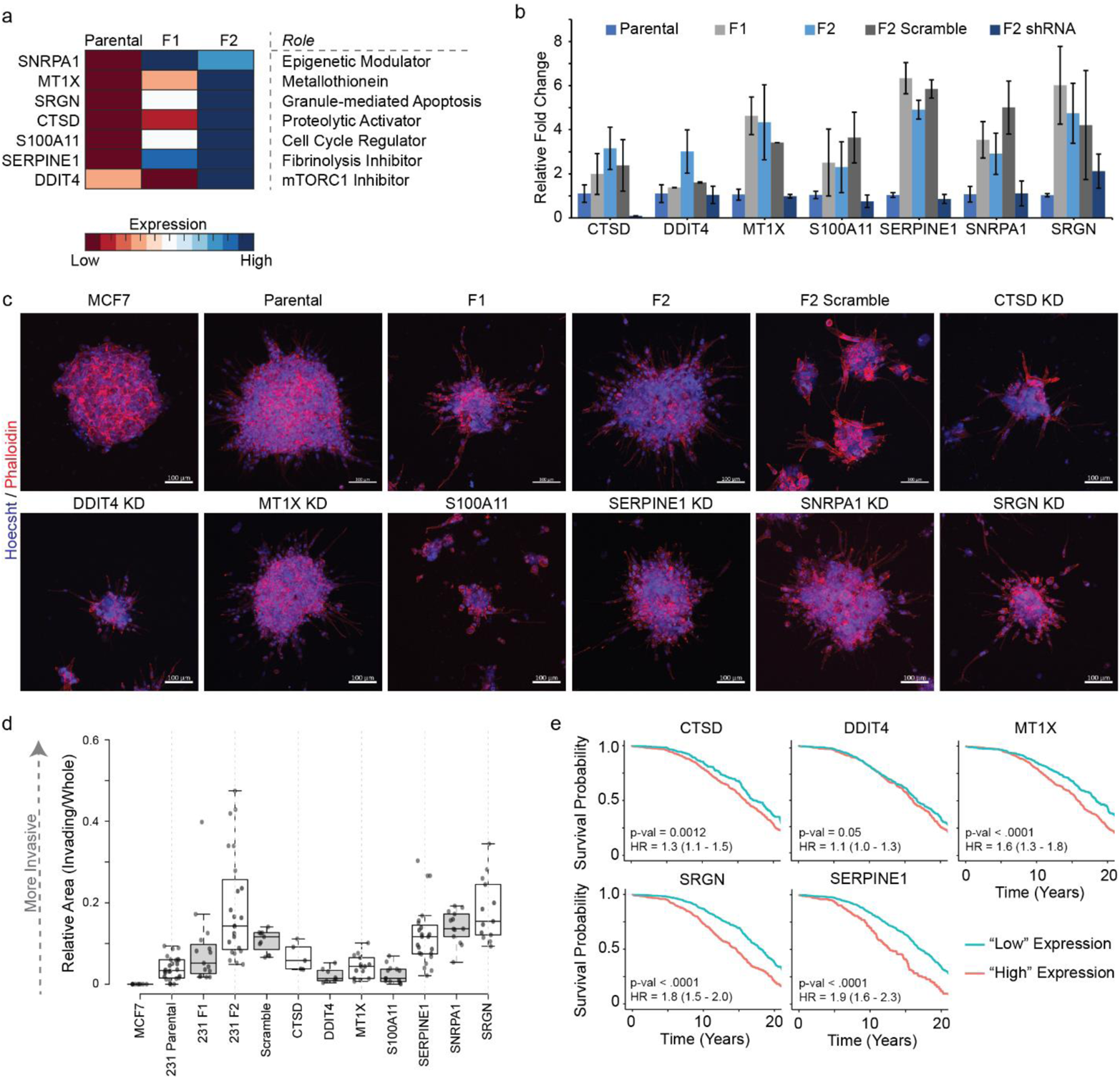
A functional assay evaluating potential transcriptomic drivers of invasion was performed. (a) Seven target genes, each with different cellular functions, were targeted for shRNA-knockdown in the F2 population. (b) qRT-PCR amplification quantifying knockdown of the respective gene within the F2 population. Notably, expression of all genes was successfully reduced to near parental expression levels. (c) Clusters of various wildtype or shRNA-knockdown cell lines were embedded within a 3-dimensional extracellular matrix and allowed to invade for 24 hours. Clusters were subsequently stained with Hoechst (blue) and Phalloidin (red) and (d) quantified for cellular invasion via ImageJ analysis. (e) Kaplan-meier survival curves analyzing gene expression with overall patient survival using the METABRIC dataset revealed statistically significant associations in 5/7 genes targeted. Scale bars = 100 µm

Next, we embedded clusters of each cell line into a 3-dimensional ECM analog and evaluated their invasion after a 24-hour period (**Fig. 4c**). Within each cluster, cells were deemed to be invading if a cellular body was identified distinct from the primary cluster body using Hoechst staining. Clusters of F2 cells infected with a scramble shRNA depicted similar invasion to wild-type F2 (*p-value* = 0.426) and F1 (*p-value* = 0.098) clusters, and statistically increased invasion relative to the MCF7 (*p-value* = 0.022) and parental (*p-value* = 0.003) clusters, consistent with their derived zebrafish phenotypes (**Fig. 4d**). Notably, shRNA-mediated knockdown of DDIT4 (*p-value* = 0.045), MT1X (*p-value* = 0.026), and S100A11 (*p-value* = 0.013) generated statistically significant decreases in invasion compared to the F2-scramble infected cells while SRGN (*p-value* = 0.473), SNRPA1 (*p-value* = 0.2954), and SERPINE1 (*p-value =* 0.152) did not decrease the invasion (**Fig. 4d**).

Finally, we sought to determine whether the genes we targeted exhibited clinical relevance within the context of metastatic breast cancer. Using publicly available clinical and gene expression data from the METABRIC study^49^, we first employed the computational tool X-tile to determine an appropriate cut-off z-score value for the genes of interest^50^. Based on this cut-off, we allocated subjects into either a high- or low-expression pool to perform Kaplan-Meier survival analysis. Of the seven genes, increased expression of all but SNRPA1 and S100A11 exhibited a statistically significant association with poorer overall survival over 240 months (**Fig. 4e**). Taken together, these data propose that DDIT4 or MT1X could be viable targets for inhibiting breast cancer invasion and metastasis.

## Discussion

Here we report the development and characterization of a model based on zebrafish xenografts that enables the rapid and robust selection of cells based on metastatic behavior. This model takes advantage of the high scalability and rapid processing potential of zebrafish embryos compared to their laboratory mouse counterparts. Through serial xenotransplantation, we generated two subpopulations of cells, F1 and F2, based solely on their ability to invade and intravasate into the zebrafish circulatory system. Previous studies have demonstrated the utility of zebrafish embryos in investigating metastasis, often stopping short of characterizing the resulting cells. Here, we have successfully enriched and expanded the subpopulation of metastatic cells from an initial heterogeneous population, resulting in cells analogous to human CTCs (**Fig. 1**). Once expanded, characterization of the populations demonstrated an ability to accurately model expected phenotypes of cancer metastasis, including an epithelial-to-mesenchymal transition and pathways involved in cancer (**Fig. 2**). In addition to transcriptomic analysis, differential RNA-splicing and protein-protein interactions in the serially generated F1 and F2 populations were consistent with expectations of invasive phenotypes from past studies (**Fig. 3**). Furthermore, functional studies involving shRNA-mediated knockdown of select genes enriched in the F2 population demonstrated a potential pipeline that could be used to identify drivers of cancer metastasis and evaluate the mechanisms of metastasis at high resolution (**Fig. 4**).

By RNA-sequencing, the F1 and F2 populations exhibited increased expression of genes that are known to be associated with cancer metastasis. For example, CSF3 (AKA granulocyte colony-stimulating factor AKA G-CSF) has been shown to act through multiple methods to enhance breast cancer metastasis, such as the induction of granulocytic myeloid-derived suppressor cells which subsequently reduce T cell activation and proliferation or through the direct activation of H-Ras oncogene, MAPK, ERK1/2, and AKT signaling pathways^51^. Similarly, mechanisms by which the top upregulated genes G0S2^52^, COL7A1^53^, IL16^54^, and DNER^55^ enhance breast cancer metastasis or seeding have all been independently verified, while increased expression of SAA2^56^ and C15orf48^57^ have been associated with poorer prognosis in breast cancers. On the other hand, loss of F1 and F2 downregulated genes such as SUSD2 has been linked to enhanced ovarian cancer metastasis in preclinical mouse models^58^. Furthermore, many of the signaling pathways enriched in the F1 and F2 populations have been identified in breast cancer and oftentimes lead to enhanced metastasis^59–62^. In addition, several pathways such as the PI3K/AKT, MAPK, and mTOR signaling pathways already have therapeutic compounds approved for treatment in breast cancer patients (e.g. rapamycin)^63^. Additionally, epithelial-mesenchymal transitions, which were shown to be upregulated in the serially generated F1 and F2 populations, have routinely been identified as critical to a cell’s ability to invade through the extracellular matrix and intravasate into the bloodstream^64^. Also, the enrichment of genes such as LPAR1 in the F2 population is consistent with previously shown enrichment in breast cancer cell line “trailblazer” populations^65^. The ability of the zebrafish model to identify enrichment of these gene sets further speaks to the accuracy by which this model can be used to study cancer metastasis.

Recently, attention has turned towards the role that alternative splicing—the process by which multiple functionally distinct transcripts can be encoded from a single gene—plays in breast cancer metastasis^66^. In-depth studies have identified global alternative splicing signatures associated with epithelial-mesenchymal transition^67^ and breast cancer metastases^43^, among other canonical pathways important to cancer metastasis. In this study, we identified differential alternative splicing in 526 transcripts spanning 442 unique genes. Of those genes with spliced variants were genes from the KEGG pathways in cancers gene set as well as those with strong associations to mammary neoplasms. The F1 and F2 subpopulations were also enriched for genes such as BIRC5 with known splice variants involved in breast cancer metastasis. PPIs add a further degree of -omics to study the zebrafish model. In this study, we found it important to evaluate expected tissue-specific interactions given the importance of extracellular signaling in cancer metastasis^68^. Among upregulated genes, MYC, VIM, and JUN were the most interconnected proteins. Similarly, APP, GABARAPL1, and MAP1LC3B were the most interconnected downregulated proteins. Silencing of APP has been shown to inhibit cell migration and invasion in breast cancers^69^. Similarly, increased protein activity of GABARAPL1^70, 71^ and ubiquitination of MAP1LC3B^72^ have both been implicated in autophagy-induced cell death, tumor invasion, and metastasis. Subsequent induction of MAP1LC3B by proteasome inhibitors induced significant tumor cell death, suggesting a therapeutic strategy for targeting PPIs^72^.

Ultimately, the zebrafish model utilized here fulfills a critical unmet need in the study of tumor metastasis^73–75^. Current studies involving metastasis are limited by the available *in vivo* technologies. The current gold standard to study tumor metastasis *in vivo* relies on murine models, which are resource-intensive and typically require months between quantifiable data points. This makes high-throughput studies impractical. Importantly the use of individual patient-derived metastatic cells and an *in vivo* model to guide clinical strategy is almost impossible using mice as the patient would succumb to the disease before sufficient data is generated. Zebrafish, on the other hand, are a useful oncogenic model that requires considerably less infrastructure, is extremely low-cost, and is uniquely suited for use within clinical timelines (~7-30 days to yield a stable cell line from zebrafish compared to months required for the same result in mice)^19^. Potential clinical applications of this model include predicting the risk of metastasis and selecting the drug (or drug cocktail) that elicits the greatest patient-specific response in the isolated and expanded metastatic cell populations.

Despite the advantages of the model demonstrated here, there are certain limitations to their application. One concern is that not all cells that intravasate successfully will become metastatic nodes^76, 77^. In this iteration of the model, experiments were stopped 5 days after injection due to the transient nature of the caudal venous plexus as an organ analogous to the bone marrow/fetal liver^78^. Recently, adult immunodeficient zebrafish analogous to severe combined immune deficiency (SCID) mice have been developed^79^. Future iterations of the model may seek to expand this work in adult *prkdc^-/-^, il2rga^-/-^* transgenic fish. Additionally, the cell lines used in this proof-of-principle study are conventional cancer cell lines, and therefore may not truly replicate the phenotype and behavior of patient-derived cells. Alternatively, injection of patient-derived cell samples would ensure greater adherence to expected metastatic behavior. These changes would enable the study of late-stage metastatic phenomena such as extravasation, dormancy, and metastatic outgrowth. Another confounding variable that needs to be acknowledged involves differences in physiological temperature between zebrafish, which are bred and maintained at 26-28°C, and humans^80^. For this study and others, xenografts are typically maintained at 34°C for the duration of their injection and monitoring, with no dramatic effects on zebrafish viability or xenograft behavior^27, 81^. Finally, despite lacking sites of secondary metastasis for breast cancer such as the brain, liver, and bone, adult zebrafish would allow the study of lung metastases, which is a major secondary site (~60% incidence) in breast cancer physiology^82^. All-in-all, we believe that this zebrafish model provides unique advantages that will likely facilitate their utility as a preclinical model for cancer metastasis over time.

In summary, we have established a rapid and robust method that leverages zebrafish to routinely isolate metastatically-inclined cells from a heterogeneous population of human cancerous cells. These cells can be expanded *in vitro* and subjected to several iterations of the *in vivo* selection process, allowing the enrichment of increasingly invasive cells. This platform has the potential to allow the rapid and inexpensive isolation and propagation of individual patient-derived metastatic cancer cells and advance basic and translational research in tumor metastasis. Given the clinical burden of metastasis, it is imperative that new technologies which take advantage of next-generation techniques be developed and are added to the cancer researcher’s toolbox. This proposed model does not require any specific device, nor does it rely on specific biomarkers, and therefore represents an easily accessible technology that can potentially be applied to any and all tumor types. Moving forward, we envision the model demonstrated here being used to identify genetic/molecular drivers of metastasis, guide the development of metastasis-prevention therapeutics, and predict clinical outcomes for individual patients.

## Materials and Methods

### Zebrafish husbandry, injections, and isolation

All animal procedures were conducted in accordance with NIH guidelines for the care and use of laboratory animals and approved all experimental protocols with zebrafish by the Georgetown University Institutional Animal Care and Use Committee, Protocol #2017-0078. For the evaluation of metastasis, cells were first labeled with the lipophilic dye CM-dil (Thermo Fisher, V22885) according to the manufacturer’s instructions. Zebrafish embryos were injected with 100-200 labeled tumor cells into the yolk sac at 2-day post fertilization (2dpf). *Tg(kdrl:grcfp)* zebrafish express green reef coral fluorescent protein in the vascular endothelium ^31^. Invasion into the vasculature was monitored on a day-to-day basis via an Olympus IX-71 inverted microscope. Embryos were evaluated daily for tumor cell migration and health of the embryos. On days of harvest, zebrafish was rinsed twice in autoclaved water followed by a brief exposure to ethanol to sterilize zebrafish. Zebrafish were then washed twice in primocin containing 1x PBS. Zebrafish were then incubated for 30 minutes in PBS mixed with 1x primocin, 1x plasmocin and Y-compound (1 µM). Tails and heads were cut using a scalpel and collected in cell culture medium supplemented with primocin and Y-compound (5µM). Tails were rinsed three times in PBS supplemented with primocin and Y-compound and subsequently spun in a centrifuge at 300g for 5 minutes at 4°C and digested in a mixture of 500 µl of PBS and Liberase at 37°C for 20 minutes. After 20 minutes, the digestion was stopped by adding 5 ml of complete DMEM and spun in a centrifuge at 300g for 5 minutes at 4°C and washed once more with PBS supplemented with primocin and Y-compound. Cells were subsequently plated in culture dishes with M-2D medium supplemented with primocin and plasmocin. Cell medium supplemented with primocin and plasmocin was changed daily for a week then every two days for additional two weeks.

### Cell Lines

MDA-MB-231 and MCF7 cells were ordered through ATCC. MDA-MB-231 cells and subsequent subpopulations were plated in medium composed of 3:1 (v/v) complete DMEM:F12 nutrient mix supplemented with insulin (final concentration 2.5 µg/mL), gentamicin (10 mg/mL), cholera toxin (0.05 nM), EGF (0.125 ng/mL), hydro-cortisone (25 ng/mL), adenine (25 µg/mL) and ROCK inhibitor Y-27632 (10 mM). All cultures were maintained at 37°C in 5% CO_2_ in a humidified chamber. Cells were split at 1-to-6 ratios every four to five days using Accutase.

### shRNA-mediated knockdown

shRNAs targeting the genes of interest were purchased from Sigma-Aldrich, including shRNAs targeting S100A11 (TRCN0000289926), SERPINE1 (TRCN0000370107), SRGN (TRCN0000007987), DDIT4 (TRCN0000062421), CTSD (TRCN0000003660), and MT1X (TRCN0000155121). shRNAs targeting SNRPA1 were gifted by Hani Goodarzi from the University of California, San Francisco. shRNA constructs were packaged in HEK 293T cells using FuGENE 6 Transfection Reagent (Promega E2691) according to manufacturer’s protocol. MDA-MB-231 F2 cells were then infected with shRNA packaged within lentiviruses in Opti-MEM (Invitrogen 51985034) supplemented with 8µg/mL polybrene (Millipore TR-1003-G). Once infected, cells were selected using increasing concentrations of puromycin up to 5µg/mL as needed. Once selected, cells were kept in culture medium supplemented with basal level puromycin (0.5µg/mL).

### 3-Dimensional Invasion Assay

Cells were dissociated into single cells using mechanical and enzymatic dissociation via Accutase (Innovative Cell Technologies #AT 104). Once dissociated, 1000 cells/well were pipetted within a 96-well U-bottom ultra-low attachment plate (Thermo Fisher #174925) followed by a quick centrifugation step at 1200 rpm for 4 minutes. Cells were supplemented with 250 uL of medium and allowed to cluster undisturbed for 4 days. After 4 days, individual clusters were collected and centrifuged at 1200 rpm for 4 minutes at 4°C. Once spun down, the supernatant was aspirated, and cell clusters were gently resuspended in neutralized rat tail Collagen I/BM mix (2.4 mg/mL Collagen I (Corning #354236) and 2 mg/mL Cultrex (BioTechne #3533-005-02), plated on 20 uL of a base layer of Collagen I/Cultrex, overlaied with serum free media, and allowed to invade for 24 hours. After 24 hours, embedded clusters were fixed for 1 hour at room temperature using 4% formaldehyde, followed by an additional 1 hour incubation step in 0.5% Triton X-100 diluted in PBS (v/v). Embedded clusters were stained for Phalloidin (Invitrogen, #A22283) and cell nuclei using Hoechst 33342 (Invitrogen #H3570) for 1 hour at room temperature. Immunofluorescence images were captured using a ZEISS LSM800 Laser Scanning Confocal. Finally, ImageJ analysis was used to quantify cellular invasion.

### RNA Extraction and qRT-PCR

Total RNA of all cell lines was isolated using Monarch Total RNA miniprep kits (#T2010S, New England Biolabs) according to manufacturer’s protocol. RNA was extracted and measured via Nanodrop-1000. RNA was subsequently sent for either library preparation and RNA-sequencing to Genewiz or cDNA preparation via a two-step process beginning with Lunascript RT Supermix Kit (#E3010, New England Biolabs). qRT-PCR was performed on cDNA from samples for genes of interest using the Luna Universal qPCR Master Mix (#M3003, New England Biolabs) according to manufacturer’s specifications. qPCR was performed on a BioRad CFX96 Touch Real-Time detection system with 60 seconds at 95°C followed by 40 cycles of 15 seconds at 95°C and 30 seconds with plate read followed by a melting curve analysis. GAPDH was measured as a reference gene. Primers used are provided in Table S1. The relative mRNA expression level was normalized to reference genes and determined using the 2^- ΔΔCT^ method.

### RNA-Sequencing Analysis

Library preparation and RNA-sequencing from isolated RNA samples was conducted by Genewiz, Massachusetts, USA using an Illumina sequencing platform. Only those RNA-samples that yielded a RIN score > 7.0 and sufficient RNA quantity were prepped and sequenced. Experimental design was made following consultation with Genewiz. Read files were trimmed using Trimmomatic ^83^ and aligned to the human genome (GRCh38.p13) using the STAR aligner ^84^. Aligned reads were quantified using featureCounts ^85^ and differential expression analysis was performed in R using DESeq2 ^86^. Normalized feature counts were used for Gene Set Enrichment Analysis (GSEA, Broad institute). Matlab R2021a was used to generate heatmaps and clustergrams for figures. In heatmaps, colors were scaled by row according to normalized feature counts.

### Differential RNA-splicing

Reads were aligned using STAR (v2.7.1a) to human genome (hg38) using iGenomes GTF annotations. MISO (v0.5.4) was then used to compare alternative splicing of skipped exons (paired end mode with fragment lengths of 219.4 (53.7) and 210.3 (52.0), for parental and derived lines respectively). Rstudio packages ggbio and ggplot2 were used to map differential splicing events and various other plots presented here.

### Protein-protein interaction and gene-disease network analysis

Network analysis for protein-protein interactions and gene-disease associations were performed using NetworkAnalyst and accessed at https://www.networkanalyst.ca ^87^. Briefly, gene lists were inputted into NetworkAnalyst. Network analysis was performed investigating breast mammary tissue specific protein-protein interactions with a stringent 30.0 filter and gene-disease associations. Network visualization was performed using a Force-atlas layout and customized using the built-in network visualization toolset.

### Kaplan-Meier Survival Curves and Clinical Data

Clinical and mRNA expression data from the METABRIC study were downloaded from cbioportal.org. Expression data, clinical vital status, and clinical overall survival in months were extracted using Matlab R2019b. X-tile software was provided online by the Rimm laboratory and accessed via https://medicine.yale.edu/lab/rimm/research/software/. To assess a suitable cutpoint for high/low expression data for each gene we used 700/1904 patient samples selected in unbiased fashion and used it as a “discovery cohort”. Cutpoints for each gene were determined using a dead of disease censor and 20-year cutoff for overall survival in X-tile ^50^. Kaplan-Meier analysis and plots were subsequently performed in Matlab R2019b using all 1904 patient samples and the respective cutpoint for each gene.

## Funding and Acknowledgements

This study was funded in part by a grant from the ACCRF and support from the Center for Cell Reprogramming at Georgetown University to SA. In addition, it was partially supported by an appointment to the Research Participation Program at CBER, US Food and Drug Administration, administered by the Oak Ridge Institute for Science and Education through an interagency agreement between the US Department of Energy and FDA (JRM). This research was also supported by the Animal Models Zebrafish Shared Resource of the Georgetown Lombardi Comprehensive Cancer Center (P30-CA051008) as well as partial funding from NIH R01CA218670.

## Author Contributions

**Conception and Design**: S. Agarwal, R, Schlegel;

**Development and Optimization of Methodology**: J. Xiao, A. Lekan, D, J. Wilkins, E. Glasgow, G. Pearson, A. Nasir, B. Johnson, K. Tanner

**Acquisition of Data and computational analysis**: J. Xiao, J. R. McGill, A. Nasir, B. Johnson

**Data Interpretation**: J. Xiao, S. Agarwal, R. Schlegel

**Writing of the Manuscript**: Initial draft is written by J. Xiao, S. Agarwal

**Editing of Manuscript:** All authors participated in editing the manuscript **Study Supervision**: S. Agarwal

**Data and materials availability**: RNA sequencing data in the current study have been deposited in the Gene Expression Omnibus (GEO) under accession number GSE153161.

## Supplemental Information

**Table S1.**
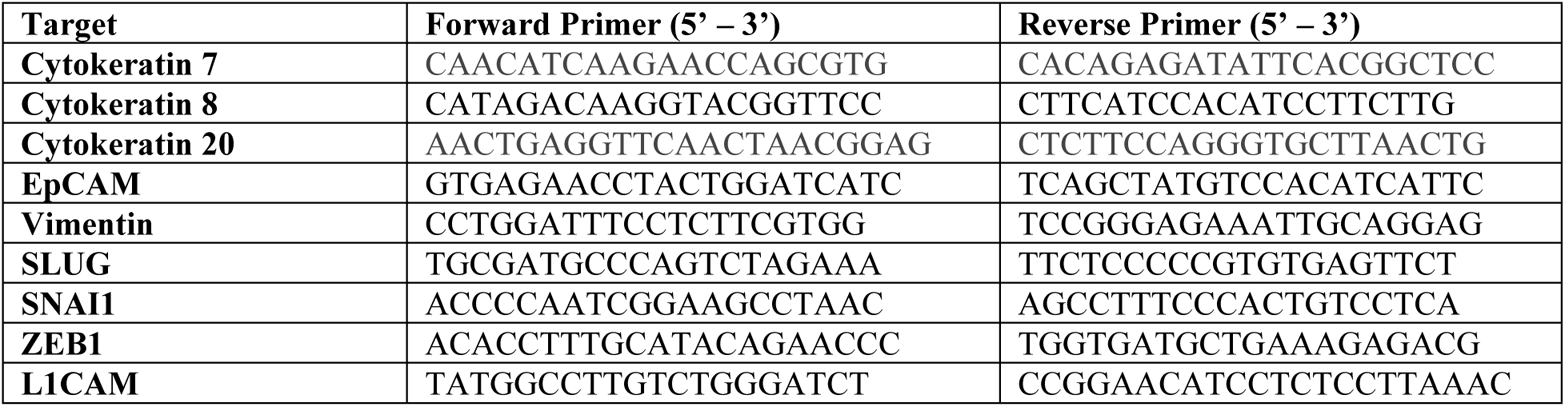
Primers used in study

**Table S2:** GSEA Enrichment using Common Parental control

| Pathway | Total | Expected | Hits | P.Value | FDR |
| --- | --- | --- | --- | --- | --- |
| TNF signaling pathway | 110 | 4.32 | 22 | 1.85E-10 | 5.90E-08 |
| Transcriptional misregulation in cancer | 186 | 7.31 | 25 | 5.75E-08 | 6.09E-06 |
| Pathways in cancer | 530 | 20.8 | 46 | 2.16E-07 | 1.49E-05 |
| NF-kappa B signaling pathway | 100 | 3.93 | 17 | 2.80E-07 | 1.49E-05 |
| MAPK signaling pathway | 295 | 11.6 | 31 | 4.50E-07 | 2.04E-05 |
| IL-17 signaling pathway | 93 | 3.65 | 16 | 5.34E-07 | 2.12E-05 |
| Cellular senescence | 160 | 6.29 | 19 | 1.49E-05 | 0.000431 |
| Cell cycle | 124 | 4.87 | 16 | 2.53E-05 | 0.000618 |
| Proteoglycans in cancer | 201 | 7.9 | 21 | 3.80E-05 | 0.000864 |
| ErbB signaling pathway | 85 | 3.34 | 12 | 0.000109 | 0.00217 |
| HIF-1 signaling pathway | 100 | 3.93 | 13 | 0.000135 | 0.00238 |
| MicroRNAs in cancer | 299 | 11.7 | 25 | 0.000277 | 0.0044 |
| Ras signaling pathway | 232 | 9.12 | 21 | 0.000296 | 0.00449 |
| Renal cell carcinoma | 69 | 2.71 | 10 | 0.000328 | 0.0046 |
| Breast cancer | 147 | 5.78 | 15 | 0.000631 | 0.00789 |
| Neurotrophin signaling pathway | 119 | 4.68 | 13 | 0.000758 | 0.0086 |
| mRNA surveillance pathway | 91 | 3.58 | 11 | 0.000823 | 0.00902 |
| Endocrine resistance | 98 | 3.85 | 11 | 0.00153 | 0.0157 |
| Rap1 signaling pathway | 206 | 8.09 | 17 | 0.00299 | 0.0265 |
| Signaling pathways regulating pluripotency of stem cells | 139 | 5.46 | 13 | 0.00313 | 0.0269 |
| Adipocytokine signaling pathway | 69 | 2.71 | 8 | 0.00538 | 0.0392 |
| PI3K-Akt signaling pathway | 354 | 13.9 | 24 | 0.00622 | 0.043 |
| mTOR signaling pathway | 153 | 6.01 | 13 | 0.00706 | 0.0468 |
| Endometrial cancer | 58 | 2.28 | 7 | 0.00727 | 0.0472 |
| Toll-like receptor signaling pathway | 104 | 4.09 | 10 | 0.00752 | 0.0478 |
| Estrogen signaling pathway | 138 | 5.42 | 12 | 0.00793 | 0.0495 |
| DNA replication | 36 | 1.41 | 5 | 0.0126 | 0.0726 |
| FoxO signaling pathway | 132 | 5.19 | 11 | 0.0146 | 0.0798 |
| Chemokine signaling pathway | 190 | 7.47 | 14 | 0.0173 | 0.0934 |
| Hippo signaling pathway | 154 | 6.05 | 12 | 0.0179 | 0.095 |
| Ubiquitin mediated proteolysis | 137 | 5.38 | 11 | 0.0188 | 0.0962 |
| Insulin signaling pathway | 137 | 5.38 | 11 | 0.0188 | 0.0962 |
| RIG-I-like receptor signaling pathway | 70 | 2.75 | 7 | 0.0195 | 0.0983 |
| VEGF signaling pathway | 59 | 2.32 | 6 | 0.0275 | 0.123 |
| GnRH signaling pathway | 93 | 3.65 | 8 | 0.0293 | 0.126 |
| EGFR tyrosine kinase inhibitor resistance | 79 | 3.1 | 7 | 0.035 | 0.145 |
| Notch signaling pathway | 48 | 1.89 | 5 | 0.0391 | 0.155 |
| Regulation of actin cytoskeleton | 214 | 8.41 | 14 | 0.0421 | 0.162 |
| Sphingolipid signaling pathway | 119 | 4.68 | 9 | 0.0442 | 0.167 |

**Table S3:** Network Nodes for Upregulated Genes

| **Label** | **Degree** | **Betweenness** |
| --- | --- | --- |
| MYC | 33 | 10770.83 |
| VIM | 21 | 8801.36 |
| JUN | 20 | 4565.89 |
| CD81 | 12 | 2763.57 |
| NFKB1 | 11 | 3404.58 |
| CEBPB | 11 | 2720.31 |
| HLA-B | 11 | 1559.45 |
| CEP250 | 10 | 3821.21 |
| EIF2C2 | 9 | 1944.19 |
| AKT1 | 9 | 1803.39 |
| TRIM28 | 8 | 5062.14 |
| KRT15 | 8 | 4475.12 |
| HSPA1A | 8 | 3307.4 |
| HDGF | 8 | 1026.54 |
| ATF3 | 8 | 578.79 |
| ALYREF | 7 | 2487.61 |
| TNFRSF1B | 7 | 1643.67 |
| GRK5 | 7 | 1301.2 |
| IKBKB | 7 | 985.72 |
| WHSC1 | 7 | 717.73 |
| E2F1 | 6 | 2220.32 |
| RUNX2 | 6 | 1382.34 |
| LYN | 6 | 1007.3 |
| DDIT3 | 6 | 383.16 |
| PKN1 | 5 | 2489.89 |
| SIAH2 | 5 | 1718 |
| TUBGCP2 | 5 | 1066.83 |
| PA2G4 | 5 | 1063.38 |
| MAGOH | 5 | 867.98 |
| HMGA1 | 5 | 746.71 |
| TNFAIP3 | 5 | 566.61 |
| LMO2 | 4 | 3364 |
| GIT1 | 4 | 1781.4 |
| CD44 | 4 | 1750.85 |
| AXIN1 | 4 | 1698.78 |
| MAP3K3 | 4 | 1240.57 |
| TUBB6 | 4 | 1046.24 |
| TFDP1 | 4 | 991.16 |
| ADRBK1 | 4 | 989 |
| DVL1 | 4 | 909.98 |
| BCL3 | 4 | 868.49 |
| **CHFR** | 4 | 776.94 |
| **EFEMP2** | 4 | 744 |
| **EIF4A1** | 4 | 691.52 |
| **RPS20** | 4 | 689.72 |
| **MAFG** | 4 | 656.85 |
| **CTBP1** | 4 | 576.44 |
| **GADD45A** | 4 | 522.24 |
| **ATAD3A** | 4 | 503.62 |
| **PPL** | 4 | 497 |
| **TRAF3** | 4 | 428.25 |
| **KLF6** | 4 | 359.3 |
| **SUV39H1** | 4 | 263.78 |
| **TRAF1** | 4 | 235.38 |
| **HDAC9** | 4 | 163.43 |
| **FOSL1** | 4 | 138.44 |

**Table S4:**
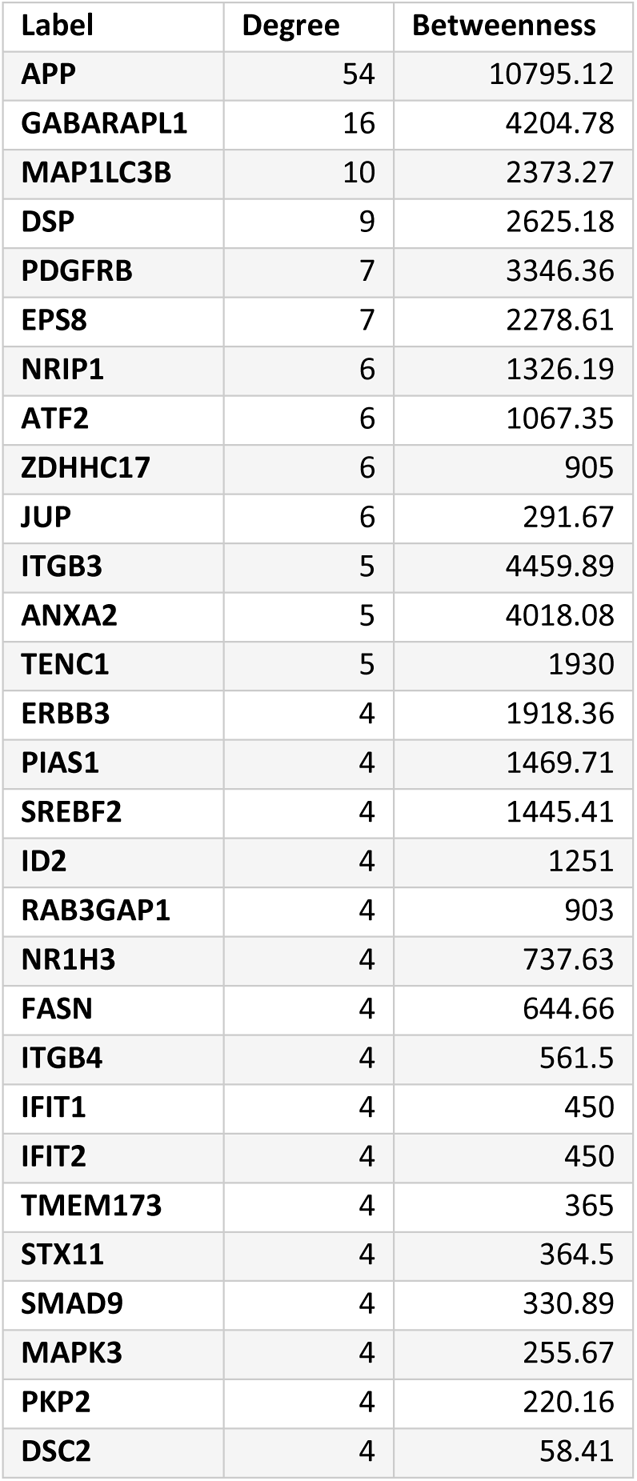
Network Nodes for Downregulated Genes, Degrees ≥ 4

